# Species-composition distributions under competition, ecological drift and immigration

**DOI:** 10.64898/2026.09.22.750472

**Authors:** Ryuichi Kumata

**Affiliations:** CEFE, Univ Montpellier, CNRS, EPHE, IRD, Montpellier, France; Institute of Tropical Medicine, Nagasaki University, Nagasaki, Japan

**Keywords:** community ecology, neutral theory, competition, coexistence, Lotka–Volterra model, demographic stochasticity, diffusion approximation, immigration

## Abstract

Understanding how species interactions, ecological drift, and immigration jointly shape community composition is a central goal of community ecology. These processes determine not only average community composition, but also how often different community states occur. Here, we ask how competition reshapes the neutral distribution of community composition generated by drift and immigration. We analyze a stochastic two-species competition model with immigration and derive a closed-form stationary distribution of relative species composition under near neutrality and weak immigration. From this distribution, we obtain explicit boundaries separating distinct unimodal, bimodal, and trimodal composition regimes. Species dominance can become common even when the corresponding deterministic model predicts stable coexistence, particularly in smaller communities, and balanced and dominance states can occur together in a trimodal distribution. Competitive asymmetry further biases composition toward the competitively favored species. Finite community size systematically shifts stochastic transitions away from their deterministic competition thresholds. Thus, competition, ecological drift, and immigration jointly determine not only how much community composition varies, but also which community states are most likely to be observed. Our results provide an analytical bridge between neutral and competition theory at the level of the full distribution of community composition.

## Introduction

Understanding how ecological processes shape community composition is a central goal of community ecology. Classical competition theory provides a deterministic framework for this question: depending on the balance between intra- and interspecific competition, two species may coexist, one may exclude the other, or alternative stable states may occur (MacArthur and Levins, 1967; Chesson, 2000; Barabás et al., 2018). Neutral theory offers a complementary probabilistic view of finite neutral communities (Hubbell, 2001; Rosindell et al., 2011). Even when species are ecologically equivalent, demographic stochasticity generates ecological drift, while immigration continually reintroduces individuals from outside the local community. Their balance determines species-abundance patterns and, in simple cases, the full distribution of community composition can be characterized analytically (Hubbell, 2001; Volkov et al., 2003; McKane et al., 2004). This raises a natural question for non-neutral communities: how does competition reshape the distribution of community composition generated by drift and immigration?

This distribution-based perspective is important because ecological processes can affect not only average composition, but also how frequently different community states occur. In natural communities, species interactions, ecological drift, and dispersal act simultaneously (Vellend, 2010; Nemergut et al., 2013; Zhou and Ning, 2017; Shoemaker et al., 2020).

Experiments show that reducing community size can increase extinction and compositional divergence even in the presence of stabilizing niche differences (Gilbert and Levine, 2017). Microbial experiments have likewise detected ecological drift directly and shown that immigration can alter compositional variation among replicate communities (Sloan et al., 2021; Catano et al., 2025). Thus, communities governed by similar ecological processes may nevertheless concentrate around very different compositions. Describing such systems therefore requires asking not only how much composition varies, but also which community states are commonly observed.

Previous stochastic competition models have shown that competition, ecological drift, and immigration can generate diverse and multimodal stationary community states (Haegeman and Loreau, 2011; Capitán et al., 2015, 2017; Lerch et al., 2023). These studies establish that stochastic community states can differ qualitatively from deterministic predictions, but the underlying composition distributions have largely been characterized numerically. As a result, we still lack a simple analytical description of how competition reshapes the distribution of relative species composition and where transitions between different distributional states occur.

Here, we analyze a minimal stochastic two-species competition model with immigration. Under weak immigration and near-neutral competition, the dynamics reduce to a one-dimensional diffusion in relative species composition, allowing its stationary distribution to be derived analytically. This provides a direct analytical link between competition, drift, and immigration and allows stochastic community states to be compared with their deterministic counterparts.

## Methods

We consider a finite local community containing two competing species and receiving immigrants from an external species pool (Fig. 1A) (Haegeman and Loreau, 2011). Let *n*_1_ and *n*_2_ denote their abundances, and let Ω set the characteristic population-size scale at which density dependence limits population growth. Individuals of both species reproduce at per-capita rate one, while immigrants of each species arrive at rate *m*Ω, so that *m* represents immigration per unit population-size scale. Competition acts through density-dependent mortality. An individual of species 1 experiences mortality proportional to *n*_1_ + *αn*_2_, whereas an individual of species 2 experiences mortality proportional to *n*_2_ + *βn*_1_. The resulting transition rates are

**Figure 1.**
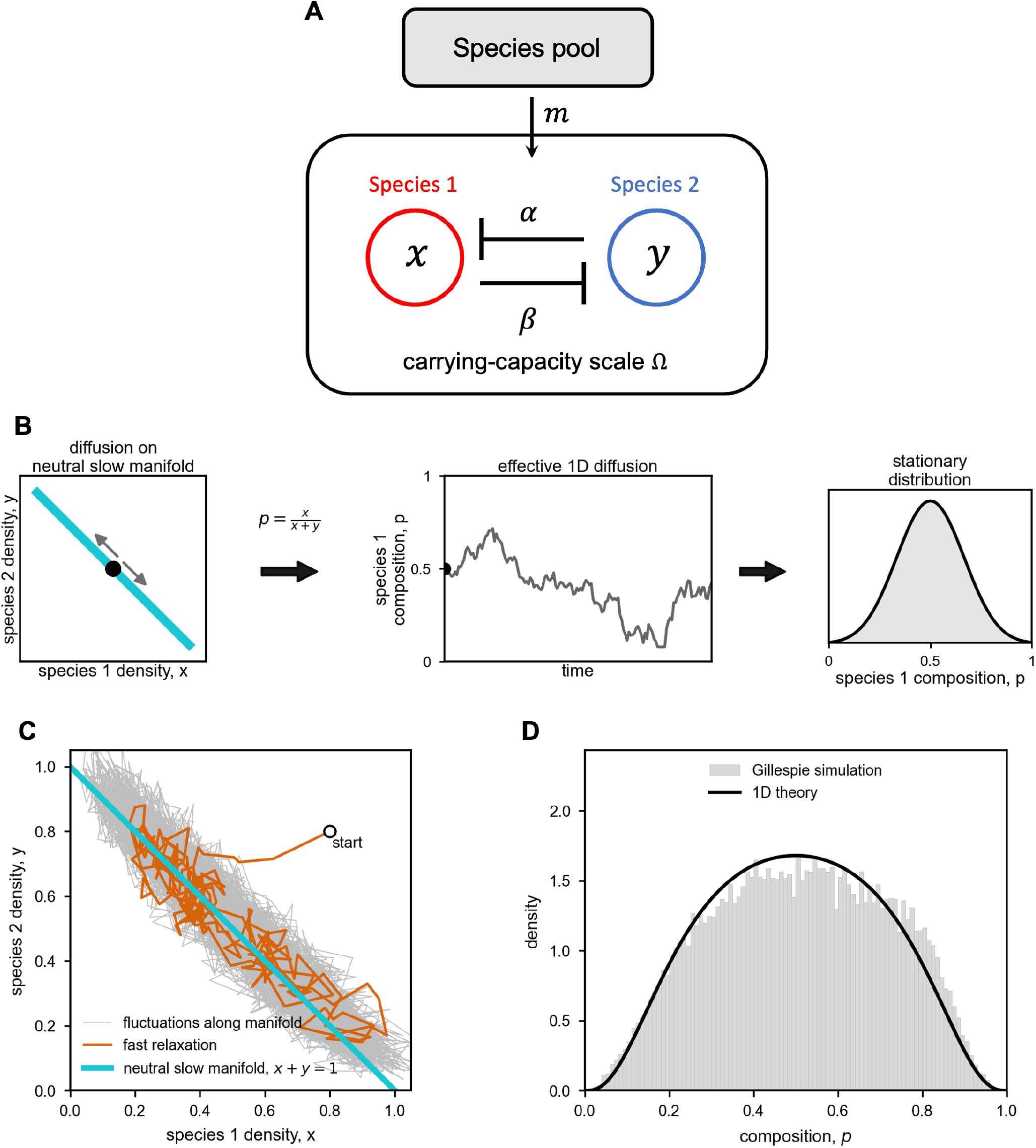
Model structure and reduction to one-dimensional diffusion. (A) Graphical representation of the competition dynamics with immigration. (B) Coordinate reduction of the two-species stochastic dynamics. Fast relaxation brings the system close to the neutral slow manifold, after which the dynamics can be approximated by one-dimensional diffusion in species composition, *p* = *x/*(*x* + *y*), leading to a stationary distribution. (C,D) Example Gillespie trajectory and stationary composition distribution. For the near-neutral parameter set shown, the trajectory rapidly approaches the vicinity of the neutral slow manifold *x* + *y* = 1 and then fluctuates mainly along it; the resulting distribution of *p* agrees with the one-dimensional diffusion theory.

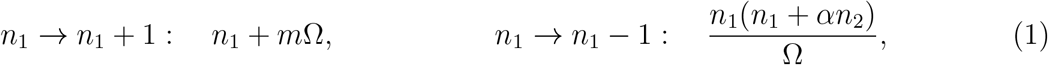

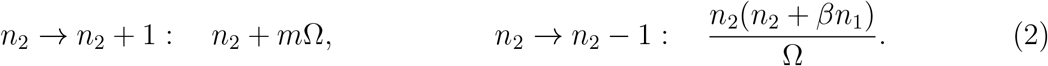

In the large-population limit (Ω *→ ∞*), these rates recover the classical two-species Lotka–Volterra competition model with symmetric immigration. Because immigration continually reintroduces both species, *n*_1_ = 0 and *n*_2_ = 0 are not absorbing states, and the stochastic system approaches a stationary distribution. Defining population densities as *x* = *n*_1_*/*Ω and *y* = *n*_2_*/*Ω, we describe community composition by the relative frequency of species 1,

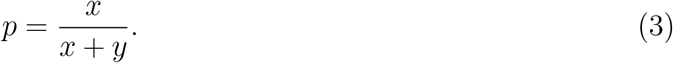

The stochastic dynamics are two-dimensional, and a closed-form stationary distribution for species composition is not generally available. For large but finite Ω, the Markov process admits a standard diffusion approximation (Gardiner, 2009). Furthermore, near neutral competition (*α ≈* 1, *β ≈* 1) and under weak immigration (*m ≪* 1), the dynamics exhibit a separation of timescales: total population density *x* + *y* relaxes rapidly, whereas relative species composition *p* changes more slowly (Fig. 1B,C) (Constable and McKane, 2015; Czuppon and Traulsen, 2018). This allows the fast density variable to be eliminated and the long-term dynamics to be reduced to a one-dimensional diffusion in species composition.

Writing *a* = *α −* 1 and *b* = *β −* 1 and eliminating the fast density variable, the following one-dimensional stochastic differential equation for composition is derived:

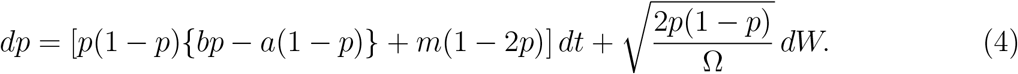

The deterministic drift term describes the effects of competition and immigration on composition, whereas the stochastic term represents demographic stochasticity, or ecological drift. Details of the reduction are given in Appendix S1.

The above reduced diffusion process of *p* has the stationary distribution

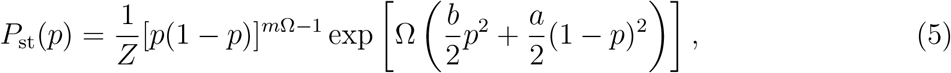

where *Z* is a normalization constant. We characterize stochastic community states by the number and location of local modes of *P*_st_(*p*), obtained from its behavior at the composition boundaries and at interior stationary points (analytical details and interpretations are given in Appendix S1). The reduced analysis accurately reproduces the stochastic trajectories and stationary distributions of the original jump process (Fig. 1C,D).

## Results

Equation (5) gives the stationary distribution of species composition within the reduced diffusion approximation. Importantly, it can be viewed as the neutral drift–immigration distribution, [*p*(1 *− p*)]^*m*Ω*−*1^, reshaped by competition through the exponential term. We characterize stochastic community states by the modality of *P*_st_(*p*), that is, by the number and location of its peaks (Fig. 2A). A mode represents a composition around which the community spends relatively large amounts of time, or equivalently a composition expected to occur frequently among replicated observations of communities under the same conditions.

**Figure 2.**
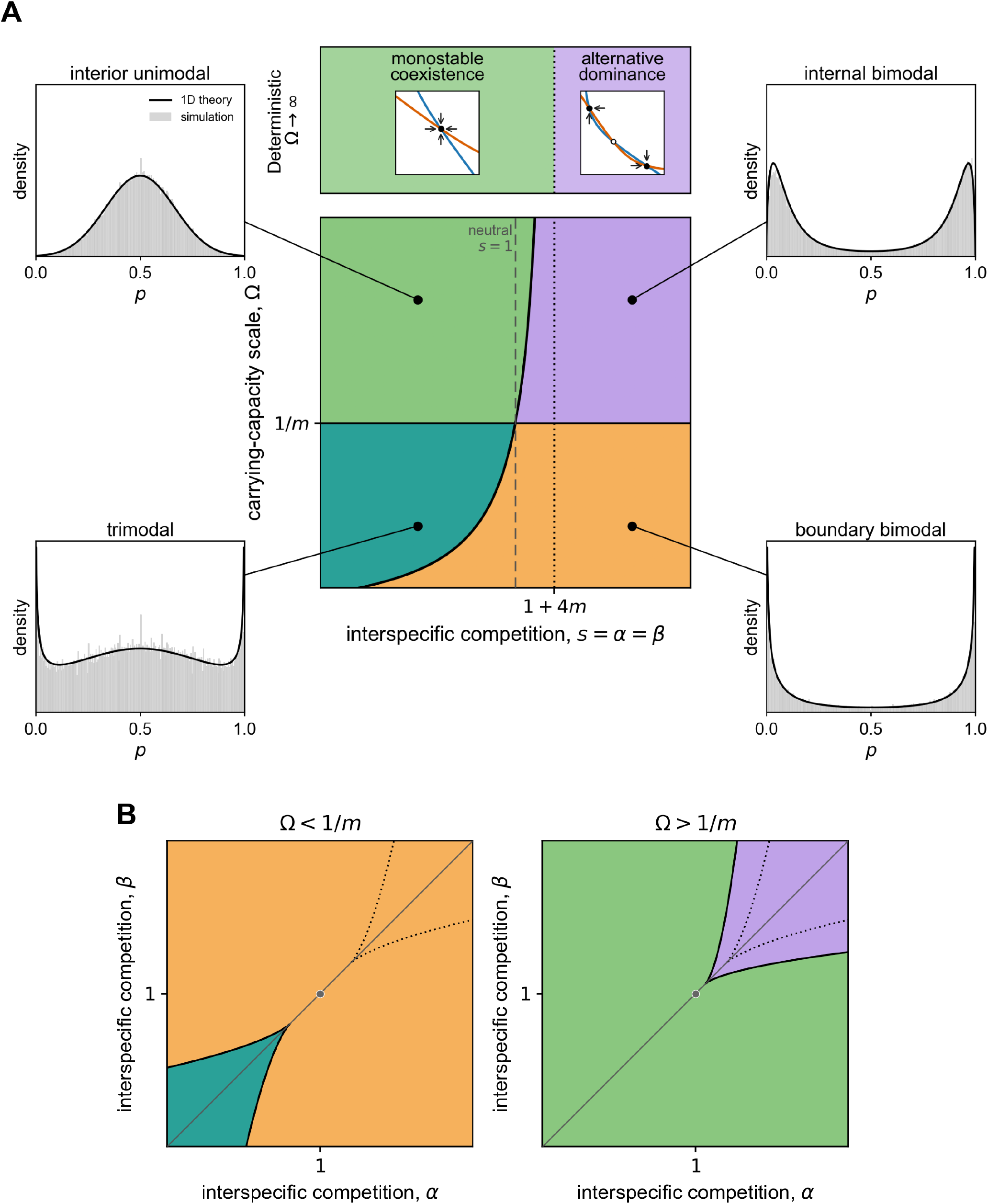
Analytical classification of species-composition distributions. (A) Phase diagram for symmetric interspecific competition, *s* = *α* = *β*, with immigration rate *m* = 0.005. Colors show the predicted modality of the stationary distribution of species 1 composition, *p* = *x/*(*x* + *y*), as Ω varies. Insets compare Gillespie simulations with the 1D diffusion theory at (*s*, Ω) = (0.95, 320), (1.06, 320), (0.95, 100), and (1.06, 100). Simulations used 100 replicate trajectories with random initial conditions, run until *t* = 8000, with a burn-in of *t* = 2000, and sampled every 5 time units. The upper band shows the deterministic limit, Ω *→ ∞*, with the transition from monostable coexistence to alternative dominance at *s* = 1 + 4*m*. (B) Corresponding (*α, β*) phase diagrams for Ω = 100 *<* 1*/m* and Ω = 300 *>* 1*/m*. Colors indicate the same stationary-distribution regimes as in panel A. Solid black curves mark stochastic modal-transition boundaries, whereas dotted black curves mark the corresponding leading-order deterministic boundaries. The gray diagonal indicates symmetric competition, *α* = *β*, and the gray point marks neutrality, *α* = *β* = 1.

Thus, an interior mode near *p* = 1*/*2 represents a commonly observed balanced composition, whereas modes near *p* = 0 or *p* = 1 represent alternative states dominated by either species. Multiple modes indicate that several distinct community compositions can be commonly observed even under identical ecological conditions.

To relate these stochastic community states to classical competition theory, we first consider symmetric competition, *α* = *β* = *s* (Fig. 2A). In the corresponding deterministic model, long-term states are described by stable equilibria: a single symmetric coexistence equilibrium is stable for

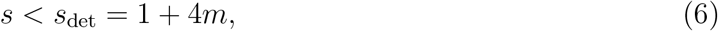

whereas two alternative stable dominance states occur for *s > s*_det_ (Fig. 2A). In finite communities, by contrast, long-term community states are described by the modes of the stationary composition distribution rather than by deterministic equilibria.

Neutral competition, *s* = 1, provides the baseline for this stochastic classification. In this case,

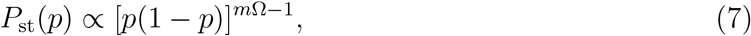

which is the symmetric beta distribution familiar from neutral community models (McKane et al., 2004). When *m*Ω *<* 1, the stationary density accumulates near *p* = 0 and *p* = 1, producing two boundary modes corresponding to alternative species dominance. At *m*Ω = 1 the distribution is uniform, whereas for *m*Ω *>* 1 it is interior unimodal and centered at *p* = 1*/*2. Thus, *m*Ω measures the balance between immigration and ecological drift, with Ω = 1*/m* defining whether boundary dominance modes are present (Fig. 2A).

Competition modifies this neutral immigration–drift baseline by determining whether the central composition *p* = 1*/*2 is a local maximum or minimum of the stationary distribution. The curvature at *p* = 1*/*2 changes sign at

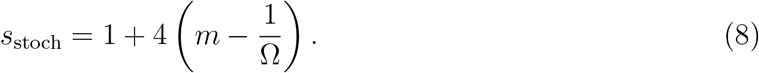

Together, the boundaries *m*Ω = 1 and *s* = *s*_stoch_ completely classify the stationary composition distributions (Fig. 2A). For *m*Ω *>* 1, the distribution is interior unimodal below *s*_stoch_ and internally bimodal above it. For *m*Ω *<* 1, the two boundary modes persist due to strong ecological drift. In this regime, sufficiently weak interspecific competition additionally produces a central mode (*s < s*_stoch_), resulting in a trimodal distribution, whereas stronger competition produces a boundary-bimodal distribution (*s > s*_stoch_).

Importantly, ecological drift shifts the loss of the central coexistence mode toward weaker interspecific competition than predicted by the deterministic model. The stochastic transition satisfies

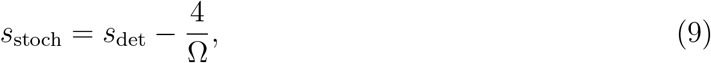

so that it shifts progressively toward weaker competition as community size decreases. Smaller communities therefore require weaker interspecific competition to maintain a central coexistence mode. Only as Ω *→ ∞* does the stochastic transition converge to the deterministic threshold.

The same classification extends to asymmetric competition (*α*≠*β*; Fig. 2B). Competitive asymmetry biases composition toward the favored species and shifts the boundaries between modal regimes. The threshold *m*Ω = 1 remains unchanged: below this threshold, species-dominance modes persist at the composition boundaries, whereas above it the modes are confined to interior compositions. Across both regimes, the stochastic modal boundaries remain distinct from the corresponding deterministic boundaries.

## Discussion

Our results show how competition reshapes the neutral distribution of community composition generated by ecological drift and immigration. Two effects organize this distribution. The balance between immigration and drift, measured by *m*Ω, determines whether species-dominance modes accumulate near the composition boundaries, whereas competition determines the location and number of interior modes. As a result, stochastic community states need not correspond directly to the equilibria predicted by deterministic competition theory. Decreasing community size shifts dominance toward weaker interspecific competition and can allow balanced and dominance states to coexist in a trimodal distribution. Competitive asymmetry further biases these states toward the favored species.

The full composition distribution contains information that is lost in summary statistics. For example, under symmetric competition, an interior-unimodal, internally bimodal, trimodal, or boundary-bimodal distribution can all have the same mean composition, *E*[*p*] = 1*/*2. Yet these distributions describe qualitatively different communities: replicates may cluster around equal abundance, separate into alternative species-dominance states, or occupy all three states. Likewise, quantities such as compositional variance, the probability that one species becomes rare, and expected diversity can all be derived from the same stationary distribution. This distributional perspective is closely related to neutral community theory, in which stochastic birth, death, and immigration generate probability distributions of species abundances (Hubbell, 2001; Volkov et al., 2003; McKane et al., 2004). Here, competition deforms this neutral distribution and connects such a probabilistic description of finite communities to classical competition theory.

Previous stochastic competition models have shown that demographic stochasticity and immigration can generate complex and multimodal abundance patterns (Haegeman and Loreau, 2011; Capitán et al., 2015; Leibovich et al., 2022; Lerch et al., 2023). Near-neutral stochastic Lotka–Volterra systems can also be reduced to lower-dimensional diffusion processes, although such reductions have been used mainly to study fixation probabilities and fixation times (Constable and McKane, 2015, 2017; Czuppon and Traulsen, 2018). By applying this reduction to the stationary distribution of relative species composition, we obtain a simple analytical classification of community states and can compare their modal transitions directly with deterministic competition boundaries. The resulting mismatch shows that deterministic stability and the long-term frequency with which different community compositions are observed are distinct properties of an ecological system.

A growing body of empirical work shows that community size and immigration can strongly influence realized community composition even in non-neutral communities. Across plant, animal, and microbial systems, smaller population or community size has been associated with stronger ecological drift, greater compositional divergence, and altered competitive outcomes even in non-neutral communities (Gilbert and Levine, 2017; Legault et al., 2019; Siqueira et al., 2020). Similarly, smaller founding populations caused highly replicated microbial communities to diverge more strongly into alternative compositional trajectories (Hayashi et al., 2026). Although founding stochasticity differs from the continual demographic drift modeled here, these studies collectively show that population size can qualitatively alter realized community states. Immigration can similarly shift the balance between stochastic and deterministic assembly. Increasing dispersal reduced compositional divergence and favored the superior competitor in experimental plant metacommunities (Ron et al., 2018), while experimental increases in immigration promoted compositional convergence in host-associated bacterial communities (Catano et al., 2025). Together, these studies show that community size, immigration, and non-neutral ecological interactions jointly affect the repeatability of community composition. Our theory suggests that such changes in observed compositional variability can reflect qualitative changes in the underlying distribution of community states. Thus, increasing ecological drift may not simply broaden variation around a typical composition, but can shift communities toward alternative dominance states or generate the coexistence of dominance and intermediate-composition states.

These results also caution against interpreting multimodal composition alone as evidence for deterministic priority effects. Priority effects arise when the order or timing of species arrival changes subsequent community assembly and can generate alternative stable or transient states (Fukami and Nakajima, 2011; Fukami, 2015; Debray et al., 2022). Empirically, however, two clusters of replicate communities dominated by different species could appear consistent with such historical contingency even when arrival history has not been manipulated. Our results show that ecological drift can produce the same qualitative pattern while the underlying deterministic system still has a single stable coexistence equilibrium. Trimodality provides an even more striking example: coexistence-like communities and communities dominated by either species can all be common under the same interaction parameters without requiring three deterministic attractors. Thus, observed alternative dominance should not by itself be taken as evidence for alternative stable states or priority effects.

The theory suggests direct empirical tests of these alternatives. If alternative dominance is strongly promoted by ecological drift, increasing community size while holding species interactions approximately constant should weaken multimodality and move replicate communities toward the central composition mode. Increasing immigration should likewise suppress accumulation near the composition boundaries, with the neutral transition determined by the product *m*Ω. In contrast, deterministic alternative stable states should not disappear solely because demographic fluctuations are reduced, and priority effects can be tested directly by manipulating species arrival order or initial composition. Experiments combining many replicate communities with factorial manipulations of community size, immigration, and assembly history would therefore allow deterministic priority effects to be distinguished from stochastic alternative dominance. Such replicated designs are becoming increasingly feasible, particularly in microbial systems (Hayashi et al., 2026; Pascual-García et al., 2025).

Our model makes several simplifying assumptions. We focused on competition close to neutrality, symmetric carrying capacities, and symmetric immigration, conditions under which the dynamics rapidly approach the nearly neutral manifold and can be reduced analytically to a one-dimensional diffusion. More strongly non-neutral interactions or demographic asymmetries are expected to deform this geometry, potentially producing a curved slow manifold. Recent projection-based diffusion methods may allow us to extend this framework to more strongly non-neutral communities by constructing one-dimensional stochastic dynamics directly along nonlinear deterministic manifolds (Sakamoto and Yeaman, 2026). A further natural extension is to multispecies systems. The neutral stationary distribution of composition is given by the Dirichlet distribution (McKane et al., 2004). Understanding how non-neutral interactions deform this neutral multispecies distribution is substantially more challenging, but represents an important direction for future work. More generally, our results suggest that community-composition distributions provide a useful complement to deterministic coexistence theory whenever ecological drift and immigration are important.

## Acknowledgments

The author thanks M. Yamamichi for insightful discussion. The author use ChatGPT (OpenAI) for assistance to revise and proofreading the draft and numerical coding.

## References

Barabás, G., R. D’Andrea, and S. M. Stump (2018). Chesson’s coexistence theory. Ecological monographs 88 (3), 277–303.

Capitán, J. A., S. Cuenda, and D. Alonso (2015). How similar can co-occurring species be in the presence of competition and ecological drift? Journal of the Royal Society Interface 12 (110), 20150604.

Capitán, J. A., S. Cuenda, and D. Alonso (2017). Stochastic competitive exclusion leads to a cascade of species extinctions. Journal of Theoretical Biology 419, 137–151.

Catano, C. P., J. G. DuBose, L. Fuller-Hall, J. Chavez, and J. C. de Roode (2025). Experimental immigration mediates ecological selection and drift in monarch microbiome assembly. Ecology Letters 28 (11), e70252.

Chesson, P. (2000). Mechanisms of maintenance of species diversity. Annual Review of Ecology and Systematics 31, 343–366.

Constable, G. W. and A. J. McKane (2015). Models of genetic drift as limiting forms of the lotka-volterra competition model. Physical review letters 114 (3), 038101.

Constable, G. W. and A. J. McKane (2017). Mapping of the stochastic lotka-volterra model to models of population genetics and game theory. Physical Review E 96 (2), 022416.

Czuppon, P. and A. Traulsen (2018). Fixation probabilities in populations under demographic fluctuations. Journal of mathematical biology 77 (4), 1233–1277.

Debray, R., R. A. Herbert, A. L. Jaffe, A. Crits-Christoph, M. E. Power, and B. Koskella (2022). Priority effects in microbiome assembly. Nature reviews microbiology 20 (2), 109–121.

Fukami, T. (2015). Historical contingency in community assembly: integrating niches, species pools, and priority effects. Annual review of ecology, evolution, and systematics 46 (1), 1–23.

Fukami, T. and M. Nakajima (2011). Community assembly: alternative stable states or alternative transient states? Ecology letters 14 (10), 973–984.

Gardiner, C. (2009). Stochastic methods (4 ed.). Springer Berlin Heidelberg.

Gilbert, B. and J. M. Levine (2017). Ecological drift and the distribution of species diversity. Proceedings of the Royal Society B: Biological Sciences 284 (1855), 20170507.

Haegeman, B. and M. Loreau (2011). A mathematical synthesis of niche and neutral theories in community ecology. Journal of theoretical biology 269 (1), 150–165.

Hayashi, I., M. Sánchez-Pinillos, and H. Toju (2026). Stochastic forces in microbial community assembly: Founding community size governs divergent ecological trajectories. Ecology Letters 29 (5), e70388.

Hubbell, S. P. (2001). The Unified Neutral Theory of Biodiversity and Biogeography, Volume 32 of Monographs in Population Biology. Princeton, New Jersey: Princeton University Press.

Legault, G., J. W. Fox, and B. A. Melbourne (2019). Demographic stochasticity alters expected outcomes in experimental and simulated non-neutral communities. Oikos 128 (12), 1704–1715.

Leibovich, N., J. Rothschild, S. Goyal, and A. Zilman (2022). Phenomenology and dynamics of competitive ecosystems beyond the niche-neutral regimes. Proceedings of the National Academy of Sciences 119 (43), e2204394119.

Lerch, B. A., A. Rudrapatna, N. Rabi, J. Wickman, T. Koffel, and C. A. Klausmeier (2023). Connecting local and regional scales with stochastic metacommunity models: Competition, ecological drift, and dispersal. Ecological Monographs 93 (4), e1591.

MacArthur, R. and R. Levins (1967). The limiting similarity, convergence, and divergence of coexisting species. The American Naturalist 101 (921), 377–385.

McKane, A. J., D. Alonso, and R. V. Solé (2004). Analytic solution of hubbell’s model of local community dynamics. Theoretical Population Biology 65 (1), 67–73.

Nemergut, D. R., S. K. Schmidt, T. Fukami, S. P. O’Neill, T. M. Bilinski, L. F. Stanish, J. E. Knelman, J. L. Darcy, R. C. Lynch, P. Wickey, et al. (2013). Patterns and processes of microbial community assembly. Microbiology and Molecular Biology Reviews 77 (3), 342–356.

Pascual-García, A., D. W. Rivett, M. L. Jones, and T. Bell (2025). Replicating community dynamics reveals how initial composition shapes the functional outcomes of bacterial communities. Nature Communications 16 (1), 3002.

Ron, R., O. Fragman-Sapir, and R. Kadmon (2018). Dispersal increases ecological selection by increasing effective community size. Proceedings of the National Academy of Sciences 115 (44), 11280–11285.

Rosindell, J., S. P. Hubbell, and R. S. Etienne (2011). The unified neutral theory of biodiversity and biogeography at age ten. Trends in ecology & evolution 26 (7), 340–348.

Sakamoto, T. and S. Yeaman (2026). Maintenance of polymorphism in spatially heterogeneous environments. Genetics 232 (1), iyaf229.

Shoemaker, L. G., L. L. Sullivan, I. Donohue, J. S. Cabral, R. J. Williams, M. M. Mayfield, J. M. Chase, C. Chu, W. S. Harpole, A. Huth, et al. (2020). Integrating the underlying structure of stochasticity into community ecology. Ecology 101 (2), e02922.

Siqueira, T., V. S. Saito, L. M. Bini, A. S. Melo, D. K. Petsch, V. L. Landeiro, K. T. Tolonen, J. Jyrkänkallio-Mikkola, J. Soininen, and J. Heino (2020). Community size can affect the signals of ecological drift and niche selection on biodiversity. Ecology 101 (6), e03014.

Sloan, W. T., C. F. Nnaji, M. Lunn, T. P. Curtis, S. D. Colloms, J. M. Couto, A. J. Pinto, S. Connelly, and S. J. Rosser (2021). Drift dynamics in microbial communities and the effective community size. Environmental Microbiology 23 (5), 2473–2483.

Vellend, M. (2010). Conceptual synthesis in community ecology. The Quarterly review of biology 85 (2), 183–206.

Volkov, I., J. R. Banavar, S. P. Hubbell, and A. Maritan (2003). Neutral theory and relative species abundance in ecology. Nature 424 (6952), 1035–1037.

Zhou, J. and D. Ning (2017). Stochastic community assembly: does it matter in microbial ecology? Microbiology and molecular biology reviews 81 (4), e00002–17.

