## supplementary appendix for "Species-composition distributions under competition, ecological drift and immigration"

### Appendix S1 for "Species-composition distributions under competition, ecological drift and immigration"

Ryuichi Kumata

**2 Stationary distribution of species composition** **5**

**3 Analytical classification of stochastic composition states** **5**

**4 Comparison with the deterministic transition** **6**

**5 Stochastic simulations** **7**

### 1 Derivation of the composition diffusion

#### 1.1 Model scaling and assumptions

We first clarify the scaling underlying the stochastic competition model. Following stochastic Lotka–Volterra formulations that combine density-dependent competition, demographic stochasticity, and immigration (Haegeman and Loreau, 2011), we consider two competing species with the same intrinsic growth rate $r$ and the same single-species carrying-capacity scale $\Omega$. We use the standard system-size scaling employed in stochastic Lotka–Volterra models (Constable and McKane, 2015; Czuppon and Traulsen, 2018). Let $N_1$ and $N_2$ denote their abundances, and let immigrants of each species arrive at rate $mr\Omega$. The corresponding deterministic dynamics are

$$\frac{dN_1}{dT} = rN_1\left(1 - \frac{N_1 + \alpha N_2}{\Omega}\right) + mr\Omega,$$
$$\frac{dN_2}{dT} = rN_2\left(1 - \frac{N_2 + \beta N_1}{\Omega}\right) + mr\Omega.$$

Here, $\alpha$ is the competitive effect of species 2 on species 1, whereas $\beta$ is the competitive effect of species 1 on species 2.

Scaling time as $t = rT$ and population densities as

$$x = \frac{N_1}{\Omega}, \qquad y = \frac{N_2}{\Omega},$$

gives

$$\frac{dx}{dt} = x(1 - x - \alpha y) + m,$$
$$\frac{dy}{dt} = y(1 - y - \beta x) + m.$$

Thus, $\Omega$ sets the characteristic population-size scale, whereas $m$ measures immigration per unit population-size scale. In the absence of immigration and the competing species, $\Omega$ is the single-species carrying capacity. Total community size is not fixed at $\Omega$ but fluctuates around a density-dependent population-size scale set by $\Omega$.

**Individual-based process** The stochastic model is constructed as the corresponding continuous-time birth–death–immigration process. After rescaling time by $r$, each individual reproduces at per-capita rate one. An individual of species 1 experiences density-dependent mortality at rate $(n_1 + \alpha n_2)/\Omega$, and an individual of species 2 at rate $(n_2 + \beta n_1)/\Omega$. Immigrants of each species arrive at rate $m\Omega$. At $\alpha = \beta = 1$, the two species are therefore demographically equivalent. The transition rates of the above individual-based process are

$$n_1 \to n_1 + 1: \quad n_1 + m\Omega, \qquad n_1 \to n_1 - 1: \quad \frac{n_1(n_1 + \alpha n_2)}{\Omega},$$
$$n_2 \to n_2 + 1: \quad n_2 + m\Omega, \qquad n_2 \to n_2 - 1: \quad \frac{n_2(n_2 + \beta n_1)}{\Omega}.$$

#### 1.2 Diffusion approximation

Introducing the densities as $x = n_1/\Omega$ and $y = n_2/\Omega$, each demographic event changes the corresponding density by $1/\Omega$. For large but finite $\Omega$, a diffusion approximation can be applied to the stochastic systems to generate the approximating diffusion process (Gardiner, 2009). A diffusion approximation of the model gives drift coefficients

$$A_x = x(1 - x - \alpha y) + m,$$
$$A_y = y(1 - y - \beta x) + m,$$

and diffusion coefficients

$$B_{xx} = \frac{x + m + x(x + \alpha y)}{\Omega},$$
$$B_{yy} = \frac{y + m + y(y + \beta x)}{\Omega},$$
$$B_{xy} = 0.$$

The cross term vanishes because no elementary demographic event changes both species simultaneously.

The corresponding Itô diffusion is

$$dx = [x(1 - x - \alpha y) + m]dt + \sqrt{\frac{x + m + x(x + \alpha y)}{\Omega}}\, dW_1,$$
$$dy = [y(1 - y - \beta x) + m]dt + \sqrt{\frac{y + m + y(y + \beta x)}{\Omega}}\, dW_2,$$

where $W_1$ and $W_2$ are independent Wiener processes. Demographic fluctuations therefore have amplitude of order $\Omega^{-1/2}$.

##### 1.3 Fast–slow reduction to species composition

Near neutrality, stochastic Lotka–Volterra systems exhibit a separation between fast population-size dynamics and slower changes in relative composition, allowing the dynamics to be reduced to dynamics along a low-dimensional slow manifold (Constable and McKane, 2015, 2017; Czuppon and Traulsen, 2018). We therefore apply the same fast–slow logic to the present model with weak immigration.

We now assume weak departures from neutrality and a large but finite population-size scale,

$$|a|,\ |b|,\ m,\ \Omega^{-1} = O(\epsilon), \qquad \epsilon \ll 1.$$

Under this scaling, competition, immigration, and demographic stochasticity all affect composition on the slow timescale, while $m\Omega$ can remain of order one.

Define total population density and relative species composition as

$$z = x + y, \qquad p = \frac{x}{x + y},$$

and departures from neutral competition as

$$a = \alpha - 1, \qquad b = \beta - 1.$$

The deterministic dynamics then become

$$\frac{dz}{dt} = z(1 - z) - (a + b)z^2 p(1 - p) + 2m,$$
$$\frac{dp}{dt} = zp(1 - p)[bp - a(1 - p)] + \frac{m}{z}(1 - 2p).$$

The quantity $bp - a(1 - p)$ represents the frequency-dependent competitive difference between the two species.

**Fast–slow structure** At exact neutrality without immigration, $a = b = m = 0$, the dynamics reduce to

$$\frac{dz}{dt} = z(1 - z),$$
$$\frac{dp}{dt} = 0.$$

Thus, for positive total population size, $z$ relaxes toward $z = 1$, whereas $p$ remains unchanged. The set $z = 1$ therefore forms a line of neutral equilibria. Linearizing around an arbitrary point $(z, p) = (1, p_0)$ on this manifold gives

$$J_0 = \begin{pmatrix} -1 & 0 \\ 0 & 0 \end{pmatrix},$$

with eigenvalues

$$\lambda_{\text{fast}} = -1, \qquad \lambda_{\text{slow}} = 0.$$

The negative eigenvalue describes rapid relaxation of total density to neutral equilibria, whereas the zero eigenvalue reflects neutrality along the composition direction.

For weak departures from neutrality, the fast manifold is slightly displaced from $z = 1$. Setting the fast deterministic dynamics to zero and expanding to first order in $a$, $b$, and $m$ gives

$$z_{\text{s}}(p) = 1 + 2m - (a + b)p(1 - p) + O(\epsilon^2).$$

Hence,

$$z_{\text{s}}(p) = 1 + O(\epsilon),$$

so total density remains close to one while composition evolves more slowly along the manifold.

**Reduction to one-dimensional diffusion** To obtain the stochastic dynamics of composition, we apply Itô's formula to

$$p(x, y) = \frac{x}{x + y}.$$

The required derivatives are

$$\frac{\partial p}{\partial x} = \frac{1 - p}{z}, \qquad \frac{\partial p}{\partial y} = -\frac{p}{z},$$
$$\frac{\partial^2 p}{\partial x^2} = -\frac{2(1 - p)}{z^2}, \qquad \frac{\partial^2 p}{\partial y^2} = \frac{2p}{z^2}.$$

Because the demographic noises of the two species are independent, $B_{xy} = 0$. Itô's formula therefore gives the drift and diffusion variance of $p$ as

$$A_p = \frac{1 - p}{z} A_x - \frac{p}{z} A_y - \frac{1 - p}{z^2} B_{xx} + \frac{p}{z^2} B_{yy},$$
$$B_{pp} = \frac{(1 - p)^2}{z^2} B_{xx} + \frac{p^2}{z^2} B_{yy}.$$

Evaluating the above expressions on $z = z_{\text{s}}(p) = 1 + O(\epsilon)$ and retaining the leading-order terms gives

$$A_p = p(1 - p)[bp - a(1 - p)] + m(1 - 2p) + O(\epsilon^2),$$
$$B_{pp} = \frac{2p(1 - p)}{\Omega} + O(\epsilon^2).$$

The leading-order one-dimensional diffusion for species composition is therefore

$$dp = [p(1 - p)\{bp - a(1 - p)\} + m(1 - 2p)]\, dt + \sqrt{\frac{2p(1 - p)}{\Omega}}\, dW.$$

The first term in the deterministic drift describes non-neutral competition, the second describes symmetric immigration toward intermediate composition, and the stochastic term represents ecological drift.

#### 2 Stationary distribution of species composition

Write the reduced diffusion as

$$dp = A(p)\,dt + \sqrt{B(p)}\,dW,$$

with

$$A(p) = p(1-p)\{bp - a(1-p)\} + m(1-2p),$$
$$B(p) = \frac{2p(1-p)}{\Omega}.$$

For a one-dimensional Itô diffusion with zero stationary probability current, the stationary density has the standard form (Gardiner, 2009)

$$P_{\mathrm{st}}(p) = \frac{C}{B(p)} \exp\left[\int^{p} \frac{2A(q)}{B(q)}\,dq\right].$$

For the present model, $m > 0$ gives inward drift at both boundaries and the resulting stationary density is normalizable.

Substituting $A(p)$ and $B(p)$ gives

$$P_{\mathrm{st}}(p) = \frac{1}{Z}[p(1-p)]^{m\Omega-1} \exp\left[\Omega\left(\frac{b}{2}p^2 + \frac{a}{2}(1-p)^2\right)\right],$$

where $Z$ is a normalization constant.

Defining

$$\mu = m\Omega,$$

the stationary density near the composition boundaries behaves as

$$P_{\mathrm{st}}(p) \propto p^{\mu-1}, \qquad p \to 0,$$
$$P_{\mathrm{st}}(p) \propto (1-p)^{\mu-1}, \qquad p \to 1.$$

The competition-dependent exponential factor remains finite at both boundaries. Therefore, boundary accumulation is determined entirely by $\mu = m\Omega$, independently of $\alpha$ and $\beta$:

$$\begin{aligned} \mu < 1 &: \quad \text{the density accumulates at both boundaries,} \\ \mu = 1 &: \quad \text{the boundary singularity disappears,} \\ \mu > 1 &: \quad \text{the density vanishes at both boundaries.} \end{aligned}$$

At neutral competition, $a = b = 0$, the stationary distribution reduces to the symmetric beta distribution familiar from neutral community and population-genetic models (McKane et al., 2004; Ewens, 2004),

$$P_{\mathrm{st}}(p) \propto p^{\mu-1}(1-p)^{\mu-1},$$

which is the density of $p \sim \mathrm{Beta}(\mu, \mu)$. Thus, $m\Omega$ determines the neutral balance between immigration and ecological drift. Note that, when immigration is absent ($m = 0$), the composition boundaries are absorbing and the long-term distribution is concentrated at fixation.

#### 3 Analytical classification of stochastic composition states

We next determine when the number of modes of the stationary composition distribution changes. Because $P_{\mathrm{st}}(p) > 0$ for $0 < p < 1$, its interior extrema $p_0$ satisfy

$$\left.\frac{d}{dp} \log P_{\mathrm{st}}(p)\right|_{p=p_0} = 0.$$

Differentiating the stationary density gives

$$\frac{d}{dp}\log P_{\text{st}}(p) = (m\Omega - 1)\frac{1-2p}{p(1-p)} + \Omega\{bp - a(1-p)\}.$$

Multiplying by $p(1-p)/\Omega$, the positions of the interior extrema $p_0$ therefore satisfy

$$F(p_0) \equiv p_0(1-p_0)\{bp_0 - a(1-p_0)\} + \left(m - \frac{1}{\Omega}\right)(1-2p_0) = 0.$$

The first term describes the frequency-dependent effect of competition, whereas the second reflects the balance between immigration and ecological drift. Thus, the locations of the modes are determined by the balance between these two effects.

A modal transition occurs when the number of interior extrema changes. At such a transition, $F(p)$ develops a multiple root at a critical composition $p = p_*$, so that

$$F(p_*) = 0, \qquad F'(p_*) = 0.$$

Solving these equations for the competition coefficients yields

$$\begin{aligned} a_{\text{st}}(p_*) &= \left(m - \frac{1}{\Omega}\right)\frac{4p_*^2 - 5p_* + 2}{p_*(1-p_*)^2}, \\ b_{\text{st}}(p_*) &= \left(m - \frac{1}{\Omega}\right)\frac{4p_*^2 - 3p_* + 1}{p_*^2(1-p_*)}. \end{aligned} \tag{1}$$

As $p_*$ varies from 0 to 1, these equations trace the stochastic modal-transition boundary ($\alpha_{\text{st}}$,$\beta_{\text{st}}$) in the $(\alpha, \beta)$ plane (Fig. 2B), where $\alpha_{\text{st}} = 1 + a_{\text{st}}(p_*)$ and $\beta_{\text{st}} = 1 + b_{\text{st}}(p_*)$.

For $0 < p_* < 1$, both rational functions multiplying $m - 1/\Omega$ are positive. Consequently, the position of the entire modal-transition boundary relative to neutrality is controlled by the sign of $m - 1/\Omega$, or equivalently by whether $m\Omega$ is smaller or larger than one.

- When $m\Omega < 1$, we have $\alpha_{\text{st}} < 1$ and $\beta_{\text{st}} < 1$. The modal transition then occurs on the weak-competition side of neutrality. In this regime, ecological drift already produces boundary modes, and sufficiently weak interspecific competition additionally stabilizes an interior mode, producing a trimodal distribution.
- When $m\Omega > 1$, we have $\alpha_{\text{st}} > 1$ and $\beta_{\text{st}} > 1$. The transition therefore occurs on the strong-competition side of neutrality. Here, boundary accumulation is absent, and sufficiently strong interspecific competition splits a single interior mode into two interior modes.
- At $m\Omega = 1$, we have $\alpha_{\text{st}} = 1$ and $\beta_{\text{st}} = 1$, and the modal-transition boundary collapses to the neutral point $\alpha = \beta = 1$.

### 4 Comparison with the deterministic transition

Since the reduction to one dimensional diffusion is derived near neutrality, we compare the stochastic modal-transition boundary with the deterministic competition model at the same leading order.

On the slow manifold, $z = 1 + O(\epsilon)$, the deterministic composition dynamics are

$$\frac{dp}{dt} = G_{\text{det}}(p) + O(\epsilon^2),$$

where

$$G_{\text{det}}(p) = p(1-p)\{bp - a(1-p)\} + m(1-2p).$$

Thus, deterministic equilibria of species composition satisfy

$$G_{\text{det}}(p) = 0.$$

A deterministic transition occurs when an interior equilibrium becomes a multiple root. At a critical composition $p = p_*$, this requires

$$G_{\text{det}}(p_*) = 0, \qquad G'_{\text{det}}(p_*) = 0.$$

Solving these conditions for the competition coefficients gives

$$a_{\text{det}}(p_*) = m\,\frac{4p_*^2 - 5p_* + 2}{p_*(1-p_*)^2},$$
$$b_{\text{det}}(p_*) = m\,\frac{4p_*^2 - 3p_* + 1}{p_*^2(1-p_*)}.$$

Compared to the stochastic modal-transition boundary (1), to leading order near neutrality, the deterministic and stochastic transition boundaries have the same geometric form, with demographic stochasticity producing the simple replacement

$$m \longrightarrow m - \frac{1}{\Omega}.$$

Finite population size therefore shifts the stochastic modal-transition boundary away from the deterministic bifurcation boundary by an amount of order $\Omega^{-1}$.

For symmetric competition, $\alpha = \beta = s$, the critical composition is $p_* = 1/2$, giving

$$s_{\text{det}} = 1 + 4m,$$
$$s_{\text{stoch}} = 1 + 4m - \frac{4}{\Omega},$$

and hence

$$s_{\text{det}} - s_{\text{stoch}} = \frac{4}{\Omega}.$$

### 5 Stochastic simulations

We evaluated the diffusion approximation against stochastic simulations of the original birth–death–immigration process. Simulations were performed using the standard Gillespie direct method (Gillespie, 1977). For state $(n_1, n_2)$, the transition rates were

$$n_1 \to n_1 + 1 \quad \text{at rate } n_1 + m\Omega,$$
$$n_1 \to n_1 - 1 \quad \text{at rate } \frac{n_1(n_1 + \alpha n_2)}{\Omega},$$
$$n_2 \to n_2 + 1 \quad \text{at rate } n_2 + m\Omega,$$
$$n_2 \to n_2 - 1 \quad \text{at rate } \frac{n_2(n_2 + \beta n_1)}{\Omega}.$$

For each parameter set, we simulated $R$ independent replicate communities. Unless otherwise stated, simulations used $R = 100$ replicate trajectories with random initial conditions, final time $T_{\text{final}} = 8000$, burn-in time $T_{\text{burn}} = 2000$, and sampling interval $\Delta t = 5$. After discarding the burn-in period, species composition was recorded as

$$p = \frac{n_1}{n_1 + n_2},$$

for states with $n_1 + n_2 > 0$. Samples from all replicate simulations were pooled to estimate the stationary distribution of $p$. No symmetrization was applied to the empirical histograms.

The empirical distributions were compared with the analytical stationary density of the one-dimensional diffusion approximation,

$$P_{\text{st}}(p) = \frac{1}{Z}[p(1-p)]^{m\Omega - 1} \exp\left\{\Omega\left[\frac{b}{2}p^2 + \frac{a}{2}(1-p)^2\right]\right\}, \qquad 0 < p < 1,$$

where $a = \alpha - 1$, $b = \beta - 1$, and $Z$ is the normalization constant.

The one-dimensional approximation is expected to be most accurate when the system rapidly relaxes toward the slow manifold and then fluctuates mainly along it, which occurs for sufficiently large $\Omega$ and competition parameters close to neutrality. Comparisons with the original jump process therefore provide a direct test of the parameter range over which the reduced diffusion accurately describes stationary community composition.

For the simulation in Fig. 1, we used a single Gillespie trajectory with $\alpha = \beta = 1.04$, $m = 0.02$, $\Omega = 200$, and initial condition $(n_1, n_2) = (160, 160)$. The trajectory was run until $T_{\text{final}} = 30000$, with burn-in $T_{\text{burn}} = 5000$ and sampling interval $\Delta t = 0.5$.

For the distribution insets in Fig. 2, we used $R = 100$ independent Gillespie trajectories with random initial conditions for each parameter set. Simulations were run until $T_{\text{final}} = 8000$, with burn-in $T_{\text{burn}} = 2000$ and sampling interval $\Delta t = 5$. The four parameter sets were given in the figure caption, with $m = 0.005$ and $s = \alpha = \beta$. Empirical histograms were estimated from pooled post-burn-in samples without symmetrization.

#### References

Constable, G. W. and McKane, A. J. (2015). Models of genetic drift as limiting forms of the lotka-volterra competition model. *Physical review letters*, 114(3):038101.

Constable, G. W. and McKane, A. J. (2017). Mapping of the stochastic lotka-volterra model to models of population genetics and game theory. *Physical Review E*, 96(2):022416.

Czuppon, P. and Traulsen, A. (2018). Fixation probabilities in populations under demographic fluctuations. *Journal of mathematical biology*, 77(4):1233–1277.

Ewens, W. J. (2004). *Mathematical population genetics: theoretical introduction*, volume 27. Springer.

Gardiner, C. (2009). *Stochastic methods*. Springer Berlin Heidelberg, 4 edition.

Gillespie, D. T. (1977). Exact stochastic simulation of coupled chemical reactions. *The journal of physical chemistry*, 81(25):2340–2361.

Haegeman, B. and Loreau, M. (2011). A mathematical synthesis of niche and neutral theories in community ecology. *Journal of theoretical biology*, 269(1):150–165.

McKane, A. J., Alonso, D., and Solé, R. V. (2004). Analytic solution of hubbell's model of local community dynamics. *Theoretical Population Biology*, 65(1):67–73.
